# Patterns of sleeping site use of Guinea baboon parties (*Papio papio*)

**DOI:** 10.1101/2025.03.20.644409

**Authors:** Lisa Ohrndorf, Roger Mundry, Jörg Beckmann, Julia Fischer, Dietmar Zinner

## Abstract

Sleeping site use in primates is thought to be influenced by multiple factors, including resource availability and competition, predation risk, and the risk of parasitic infection. While previous research has highlighted the importance of these factors, few studies have examined sleeping site use among multiple groups of non-territorial species with overlapping home ranges. Here, we investigated the sleeping site use of Guinea baboons (*Papio papio*) in Simenti, Senegal. We used locational data for four years of several baboon parties sharing the same range. We assessed the distribution of sleeping sites within the local habitat mosaic, the proximity of sleeping sites to those of co-occurring parties and the impact of food availability and predator presence at the landscape level on the distance between parties’ sleeping sites on the same night. We further investigated patterns of sleeping site use of parties across time. We found that the vast majority of sleeping sites were in the gallery forest along the perennial Gambia River, likely due to the availability of tall trees. Guinea baboons slept in close spatial proximity (>50m) to at least one other party regardless of food availability and predator presence. Patterns of sleeping site use showed no evidence of predator or parasite avoidance. Thus, Guinea baboons in Simenti likely use the abundantly available sleeping sites opportunistically rather than being driven by intergroup competition or strategies for predator avoidance or reduction of the risk of parasite infection.

## Introduction

Choosing suitable and safe sleeping sites is considered critical to individual survival and fitness in many primate species (Altmann, 1974; Anderson, 1984, 1998; Reichard, 1998). Sleeping site choice is often seen as an optimisation process with which individuals or groups balance the effects of multiple variables, including resource availability and competition (Chapman, 1989), predation risk (Anderson, 1984), territory and range defence (Heymann, 1995), and exposure to parasites (Hausfater & Meade, 1982; Anderson, 1998). Depending on the ecological conditions within their habitats, primates are expected to select sleeping sites near feeding sites or water bodies, as sites closer to such resources allow individuals to spend more time feeding by minimising travel distances and associated costs between sleeping sites and feeding patches (Chapman, 1989). Such spatial relationships between essential resources were found in, for instance, ursine colobus monkeys (*Colobus vellerosus*; Albert et al., 2011), pigtailed macaques (*Macaca leonina;* Teichroeb et al., 2012), and hamadryas baboons (*Papio hamadryas*; Sigg & Stolba, 1981; Schreier & Swedell, 2008; Henriquez et al., 2021).

Another non-mutually exclusive driver of sleeping site selection may be predation risk. Many primate species are diurnal and inactive at night, making them vulnerable to nocturnal predation (Cowlishaw, 1994). Therefore, selecting safe sleeping sites may be essential for protection from predators. Various primate species prefer sleeping sites with specific attributes, such as trees that are tall, have large diameters at breast height (DBH), or difficult-to-climb trunks that likely offer better protection from nocturnal predation (e.g., titi monkeys *Callicebus coimbrai*; Souza-Alves et al., 2011); black-fronted titi monkeys (*Callicebus nigrifrons*; Caselli et al., 2017); Bornean agile gibbons (*Hylobates albibarbis*; Cheyne et al., 2012).

Besides selecting trees or tree species with such protective attributes, primates may also employ a strategy of switching between multiple sleeping sites to minimise predation risk by reducing the predictability of their nocturnal location. By having access to and switching between multiple sleeping sites within their home ranges, individuals or groups can thus decrease the likelihood of being detected by predators (Hamilton, 1982; Markham et al., 2016; Caselli et al., 2017).

However, the availability of suitable sleeping sites to switch between may vary strongly between habitats. Species inhabiting forested areas that sleep openly on the branches of tall trees might find an abundance of suitable sleeping sites to choose from in their habitat, whereas other species or populations that live in more open habitats may have access to only a few sites within their home ranges (e.g., hamadryas baboons; Kummer, 1968; Sigg & Stolba, 1981). For such species, suitable or high-quality sleeping sites are often considered a limited resource, similar to species that sleep in tree holes. A scarcity of suitable sleeping sites may lead to competition for such sites among groups or sub-groups that share a home range (Altmann, 1974; Cheyne et al., 2012; Markham et al., 2016).

Another common hypothesis is that primates switch sleeping sites to minimise exposure to faecal matter build-up, which could otherwise increase the risk of parasitic infection (Hausfater & Meade, 1982; von Hippel, 1998). Given the predictability of sleeping site use for predators and the faecal matter build-up, the quality of sleeping sites should decrease when used continuously (Hamilton, 1982; Markham et al., 2016). Patterns of sleeping site use in line with parasite avoidance have been found in yellow baboons (*P. cynocephalus*) in Amboseli, Kenya (Hausfater & Meade, 1982; Markham et al., 2016). However, the intervals of vacancies for sleeping sites of individual groups were shorter than the predicted optimum to avoid parasite contact, and the occupancy was almost constant when looking at sleeping sites shared among groups.

Examining the simultaneous use of sleeping sites by multiple groups is essential for understanding the usage patterns, particularly in non-territorial species with overlapping home ranges. In such species, sleeping sites are likely to be used by several groups or sub-groups simultaneously or successively, potentially influencing the dynamics of site selection regarding predation risk and parasite load.

In this study, we investigated patterns of sleeping site use of Guinea baboon parties co-occurring within the same range. Guinea baboons (*P. papio*) are group-living, non-territorial primates that are highly spatially tolerant (Ohrndorf et al., 2025). They form large multi-male, multi-female groups of 20 to up to more than 300 individuals that often aggregate at shared sleeping sites (Dunbar & Nathan, 1972; Sharman, 1982; Galat-Luong et al., 2006; Patzelt et al., 2011). Guinea baboon societies show a nested multi-level social organisation with fission-fusion dynamics. At the core of this social organisation is the one-male unit, consisting of one primary male, one to several females, their dependent offspring, and sometimes secondary males (Goffe et al., 2016). Several one-male units form a party, which, together with other parties, form gangs (Fischer et al., 2017). In the Niokolo-Koba National Park in Senegal, the home ranges of parties vary between 20 and 50 km^2^ (Sharman, 1982; Zinner et al., 2021) and overlap with parties belonging to the same gang almost entirely (Ohrndorf et al., 2025; Zinner et al., 2021). Guinea baboons in Simenti live in a savannah woodland mosaic, and their sleeping sites are predominantly located in the core areas of their home ranges, often in the gallery forest alongside the Gambia River. Most sleeping sites in the study area consist of one or several trees of *Ceiba pentandra*, *Celtis integrifolia*, and *Borassus akeassii* (Zinner et al., 2021), abundant in habitats bordering perennial water bodies. Like other baboon species, Guinea baboons are eclectic omnivores that primarily feed on fruit but also other plant parts, invertebrates, and occasionally small birds or mammals (Sharman, 1982; Zinner et al., 2021, O’Hearn et al., 2024). The most important predators of Guinea baboons in Simenti are African lions (*Panthera leo*), leopards (*P. pardus*), and spotted hyenas (*Crocuta crocuta*).

Using locational data from four years, we analysed the use of sleeping sites of Guinea baboon parties in Simenti, Senegal. We describe patterns of sleeping site use regarding the habitat type in which sleeping sites were located. We assessed the sleeping site use of Guinea baboon parties in relation to other parties that share the same range and in response to variations in food availability and predator presence across the landscape. Additionally, we looked at patterns of sleeping site use of individual parties to assess whether we could find any pattern indicative of predator or parasite avoidance. With this, we aim to provide additional information on the drivers of sleeping site use in Guinea baboon parties.

## Material & Methods

### Study site

The study took place at the long-term field site of the Centre de Recherche de Primatologie (CRP) in Simenti, Senegal (13°01‘34‘‘ N, 13°17‘41‘‘ W). The site is located in the Niokolo-Koba National Park in southeast Senegal and belongs to the Sudanian and Sahelo-Sudanian climatic zone (Arbonnier, 2002). The vegetation in the study area around Simenti is classified as a mosaic of grasslands, wooded savannahs, and gallery forests alongside rivers and other perennial water bodies (Arbonnier, 2002; Burgess et al., 2004). Seasonality in this climatic zone is pronounced, with an annual precipitation of 950 mm, mainly limited to the rainy season. The rainy season typically lasts from June to October, with May and mid-October constituting transitional periods with highly variable rainfall (Zinner et al., 2021). Several habituated Guinea baboon parties have been followed extensively since 2007 for behavioural studies at our study site.

### Data collection

#### GPS data

To determine the location of sleeping sites of Guinea baboon parties within the study area, we deployed GPS collars (Tellus 2 Basic Light) with built-in drop-off mechanisms on eight adult male Guinea baboons from eight different parties (P). Each male represented their entire party’s location (P5, P6I, P6W, P7, P9B, P13, P15, P17), with party membership determined through regular observations of social interactions and spatial proximity (Patzelt et al., 2014). We programmed the collars to take locational fixes at 2-hour intervals during the day (06:00-18:00) and an additional three fixes during the night (21:00, 00:00, 03:00).

To assess the location of the respective sleeping site, we subset the dataset of locational fixes to only those taken at night. From these fixes, we calculated the centre for each individual per night to estimate the location of the sleeping sites of their respective parties (Figure 1). We refer to “sleeping sites” for a party as the nocturnal locations of the collared individual rather than as discrete, identifiable sites, as delineating such sites, e.g., in continuous gallery forest, was not possible due to almost complete use of sleeping trees in the gallery forest alongside the Gambia River (Figure 2). The distance between locational fixes taken at 21:00, 00:00 and 03:00 for each party was 10 m on average (median, 6.75 – 15.25 m IQR), which can be caused by the nocturnal movements of the collared individuals or by GPS inaccuracies.

**Figure 1:**
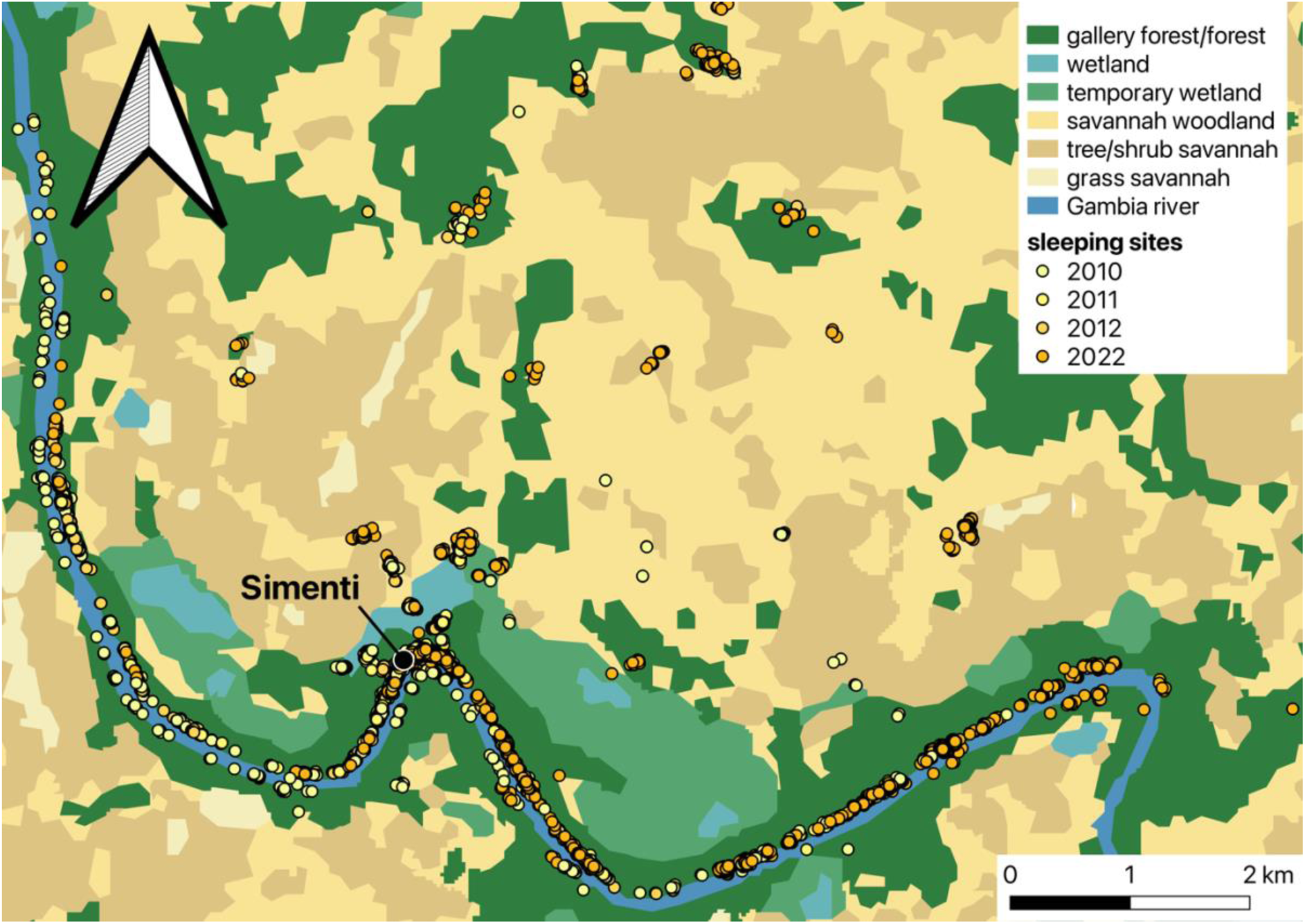
Locations of sleeping sites (spatial centre of GPS fixes taken at 21:00, 00:00 and 03:00 each night) in the different habitat types within the study area for all parties surveyed from 2010-2012 and in 2022. The true number of sleeping site locations is obscured by extensive overlap.

**Figure 2:**
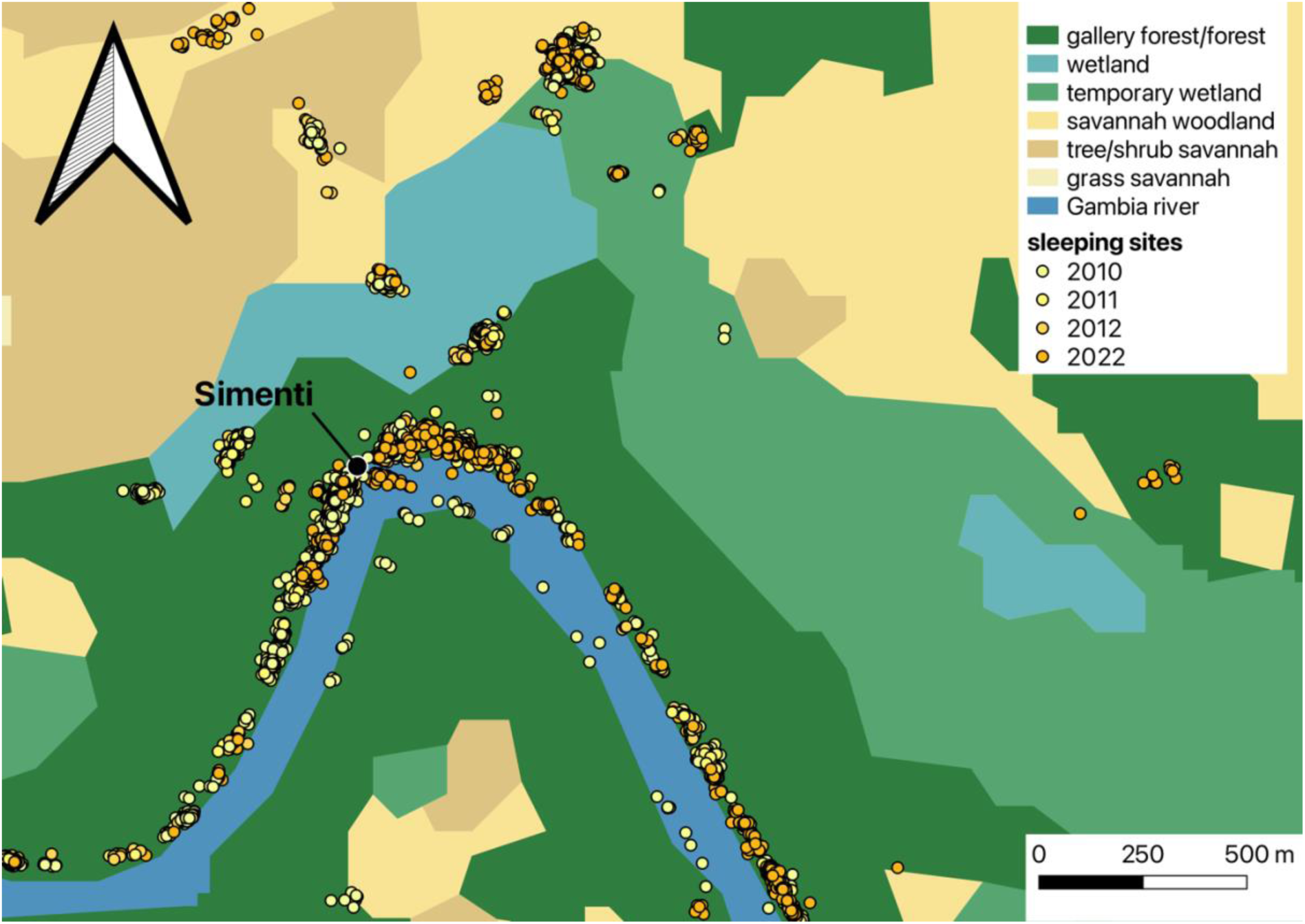
Nearly complete use of sleeping trees in the gallery forest along the Gambia River by all study parties from 2010-2012 and 2022. Sleeping sites in the water are likely due to GPS inaccuracies or overhanging branches of sleeping trees.

Additional GPS data were available from a previous study for eleven individuals (four females, seven males) from six parties in 2010, seven individuals (two females, five males) from six parties in 2011, and five individuals (one female, four males) from five parties in 2012 (Patzelt et al., 2014). The sampling intervals for these collaring periods were identical to the ones in 2022. The collaring was approved by the Ethics committee of the German Primate Center (document number E4-21). For a detailed description of the capture and collaring procedure from 2010 to 2012, as well as for 2022, please see Knauf et al., 2015 and Ohrndorf et al., 2025, respectively.

#### Habitat types

To understand in which habitat types sleeping sites of Guinea baboon parties predominantly occur, we used the habitat classification established via remote sensing of Landsat 5 TM imagery from a previous study (Klapproth, 2010) (Figure 1). We further investigated whether the use of different habitat types for sleeping sites changed within and across years.

#### Proximity among sleeping sites of parties

To assess patterns of proximity between the sleeping sites of our study parties, we assessed the minimum Euclidean distance between the sleeping sites of pairs of collared individuals, each representing their entire party’s location, for each night GPS data were available for both individuals. Then, we identified the closest neighbour of each of the eight study parties and the associated shortest distances between them until each party was represented at least once for every night they were observed simultaneously as another party (Figure 3). We chose to use minimum distances between parties instead of average distances because, in a limited space, movements away from one party would often bring individuals closer to another, leading to averages not being able to capture the spatial relationships we were interested in.

**Figure 3:**
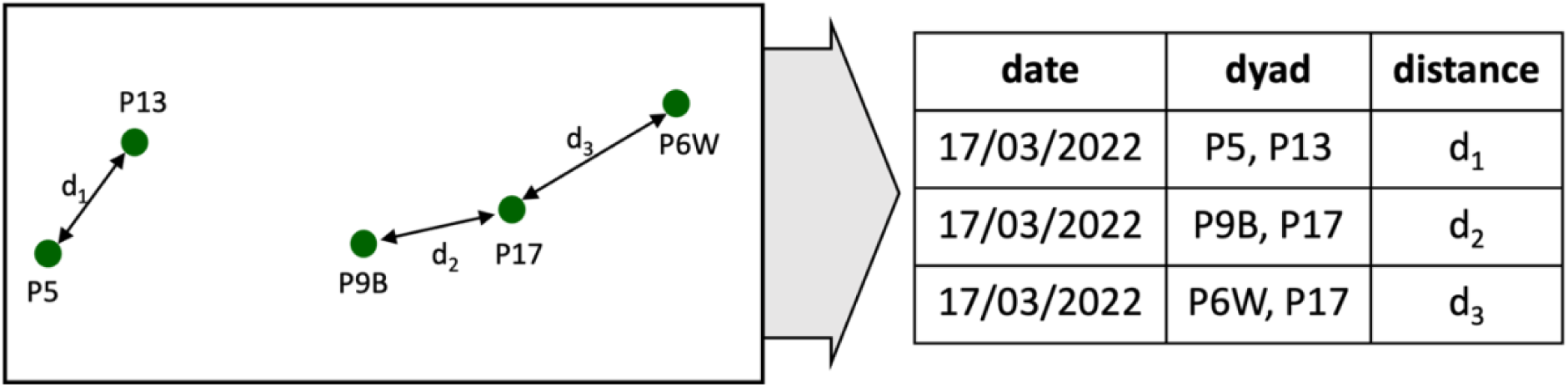
Example of the assessment of minimum distances between sleeping sites of parties. In this example, GPS data were available for five parties (P5, P6W, P9B, P13, P17). We identified the closest party dyads (P5 and P13, P9B and P17, P6W and P17) and the Euclidean distances between them (d_1_-d_3_).

#### Food availability

To evaluate food availability across our study parties’ home ranges, we used the same phenological data as described in Ohrndorf et al. (2025). We conducted monthly phenological assessments of key feeding tree species from November 2021 to March 2023. We selected an average of ten trees (range: 1-13) from each of the 31 most common feeding species identified by Zinner et al. (2021). These marked trees were monitored at the start of each month throughout the 17-month study period. We recorded eight phenological states: none, young leaves, flower buds, flowers, young fruits, intermediate fruits, ripe fruits, and mature leaves. Trees with ripe or intermediate fruits were classified as “providing food,” as were flowering trees of species known to be consumed by our study population. We then determined food availability as the average of the proportions of trees providing food per species.

#### Predator presence

We assessed predator presence across the study area using unbaited, motion-triggered camera traps distributed across 37 km^2^ as described in Ohrndorf et al. (2025), covering most of our study parties’ home ranges. We deployed the cameras on a grid of 1 x 1 km, with roughly one camera per km^2^. The imagery obtained from this camera-trapping grid was annotated using the online platform *Agouti* (Casaer et al., 2019). We checked the AI-assisted annotation manually (Brouillet, 2024). We considered sightings of lions, leopards, hyenas, and two sightings of African wild dogs (*Lycaon pictus*) within the camera-trapping grid as predator encounters.

In addition to the camera trap data, we collected ad libitum data on all signs of predators (tracks, scat, calls, sightings) during the study period. From both camera trapping and ad libitum observations, we calculated the number of predator encounters in the study area in intervals of 14 days. As we could not be certain about the most relevant time window for assessing perceived predation pressure in Guinea baboons, we also calculated the number of predator encounters over shorter (2 and 7 days) and longer (30 days) intervals.

#### Effect of food availability and predator presence

To investigate whether the distance between sleeping sites of study parties varied with food availability and predator presence, we used data on food availability and predator presence and minimum distances between sleeping sites of study parties from 2022. For the years 2010 to 2012, there were no data available on food availability or predator presence. We fitted multiple membership models using the function *brm* of the package *brms* version 2.20.4 (Bürkner, 2017) in R version 4.3.1 (R Core Team, 2023). We log-transformed the minimum distance between parties per night to increase the probability of model convergence and included the log-transformed distance as the response variable in our models. We included the monthly food availability score and the number of predator encounters within 14 days prior to the observation as predictors. To account for variation between individuals and dyads of individuals, we included the IDs of both individuals in a dyad (as a multi-membership term) and dyad ID as random intercepts effect. We also included random slopes for food availability and predator presence within the 14-day period to avoid overconfident model estimates (Barr et al., 2013; Schielzeth & Forstmeier, 2009).

The multi-membership approach estimates a single random intercepts effect for both individuals in a dyad. Thus, the dyad effect accounts for preferential associations between specific parties that may be generally closer to some parties but not others. We chose a multi-membership model as the two individuals of a dyad could not be unambiguously assigned to two different random effects variables. Before fitting the model, we checked for sufficient variation in food availability and the number of predator encounters within each individual and dyad. We also included parameters for the correlations among random intercepts and random slopes in the model. As we received a warning about divergent transitions during warm-up with the default settings of *brm*, we set adapt_delta to 0.99.

To ensure that our model results were not biased by an inappropriate choice of the time window used to assess predator presence, we fitted three additional models, including the number of predator encounters per 2, 7 and 30 days instead of 14 days as predictors, but maintaining an otherwise identical structure.

#### Patterns of sleeping site use

We visually inspected how individual parties used sleeping sites by plotting the distances between successive sleeping sites across time. We then randomised the order in which each party visited their sleeping sites 1000 times to see whether the observed distribution of distances between sleeping sites differed from the random expectation. There are four different patterns we might expect:

A. Guinea baboon parties prefer to use sleeping sites close to that used the previous night with only a few larger distances between successively used sites, indicating frequent use of a few select sites or overall mainly small scale replacement of successive sleeping sites (Figure 4A).
B. Guinea baboons might avoid detection by predators or reduce exposure to parasites from faecal matter build-up. In this case, we expect very few short distances and many medium to long distances between successive sleeping sites to minimise predictability for predators or reduce the risk of parasite infection (Figure 4B).
C. Alternatively, we might observe many short distances on days when Guinea baboons use the same sites or sites close by, combined with a considerable fraction of long distances observed when they switch to areas far from previously used ones and use sleeping sites there to avoid predator detection (or after disturbance by a predator) or contamination (Figure 4C).
D. Guinea baboons might use sleeping sites randomly, visiting sites at all distances from previously used sites with equal likelihood (Figure 4D).

**Figure 4:**
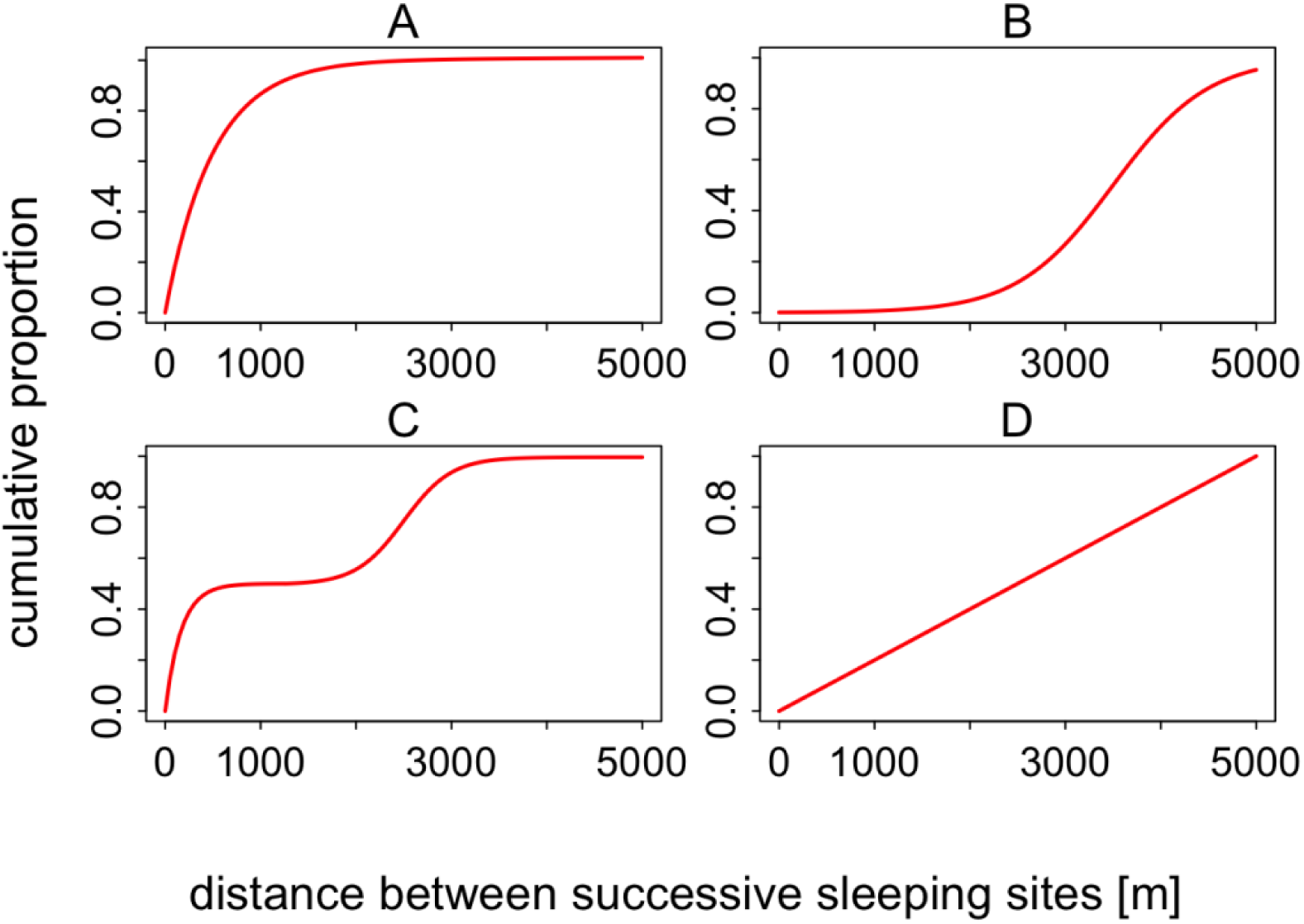
Cumulative proportion of distances between successively used sleeping sites under four hypothetical scenarios (A–D). The x-axis represents the distance between successive sleeping sites. The y-axis shows the cumulative proportion of data, indicating the percentage of the total data as distance increases.

## Results

We collected the geographical coordinates of 8913 sleeping sites across all years and parties. In 2022, GPS collars lasted an average of 316 days, resulting in 2532 sleeping site locations across eight study parties. One collar (P6I) failed after 162 days due to water damage.

### Sleeping sites and habitat types

Out of 8913 recorded sleeping sites, 7480 fell into gallery forests, 473 into wetlands, 449 into temporary wetlands, 432 into savannah woodlands, and 78 into tree/shrub savannahs. One sleeping site fell outside of the geographic range of the habitat classification Klapproth, (2010) and could, therefore, not be assessed. Across all years, our study parties spent 84% of their nights in the gallery forest and other forest types, despite these habitat types covering only 15.5% and 14.8% of the area, respectively (Klapproth, 2010) (Figure 5). Our study parties spent 5% of their nights in trees near wetlands or temporary wetlands. When looking at habitat use for sleeping site locations across the years, most sleeping sites were in the gallery forests year-round, especially during the rainy season (June – October). During the dry seasons, there was more variability in habitat use, with parties spending more nights in other habitats such as wetlands, temporary wetlands, or savannah habitats. During the dry season of 2022, our study parties spent substantially more nights in savannah woodlands than during the rainy season (Figure 5).

**Figure 5:**
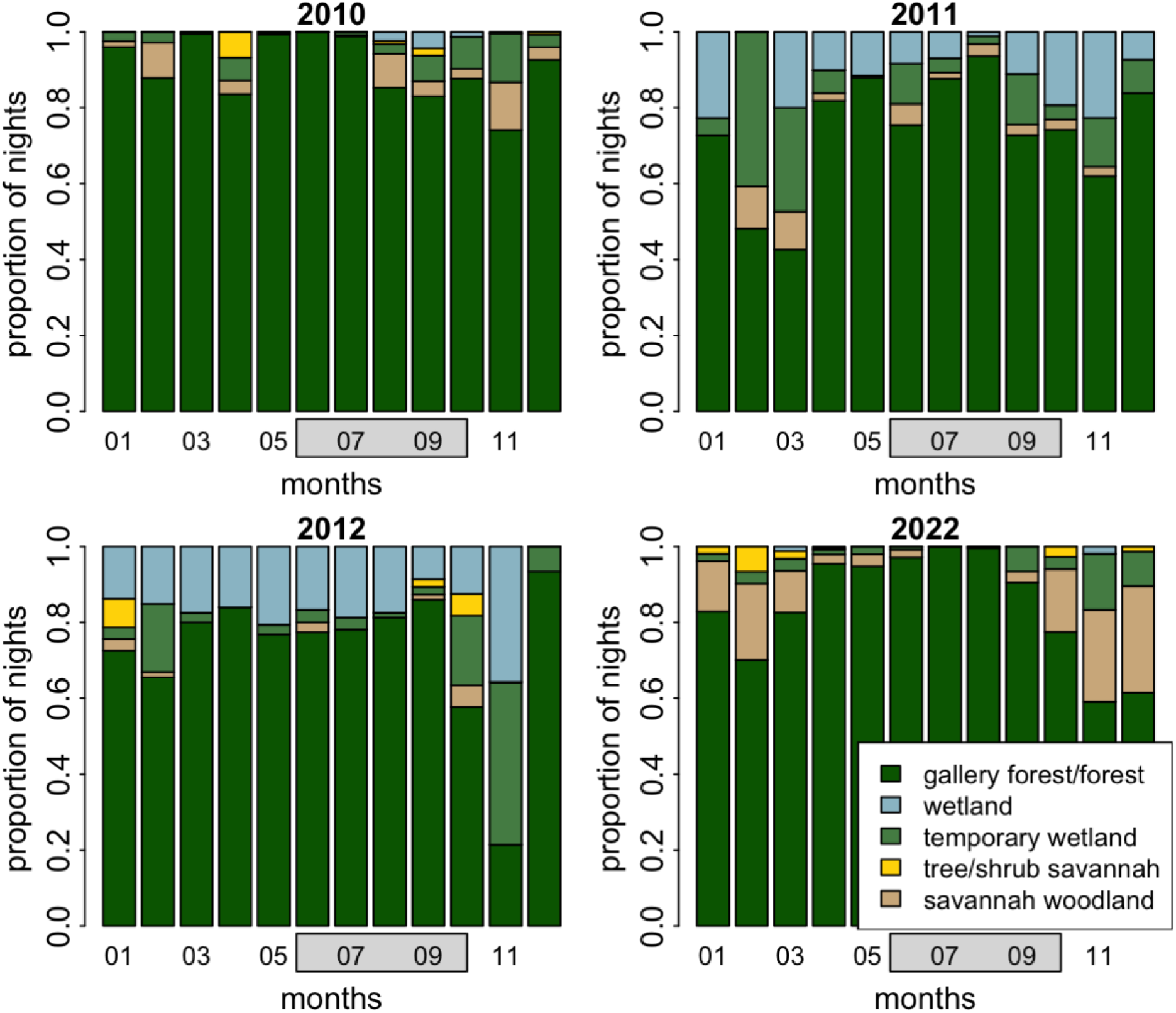
Proportion of nights spent in each habitat type per month across the years of observation. Grey boxes represent the months of the rainy season (June to mid-October).

### Food availability and predator presence

Food availability scores fluctuated between 0.03 and 0.33 across the study period (Figure 6A). The highest scores were observed in February, March, and April 2022, while the lowest scores occurred in June, July, and August, despite these months falling within the rainy season. In 2022, we recorded a total of 588 predator encounters (Figure 6B). This included 376 ad libitum records and 212 images of predators taken by camera traps. Spotted hyenas were the most frequently recorded predators (254 encounters), followed by lions (211) and leopards (118).

**Figure 6:**
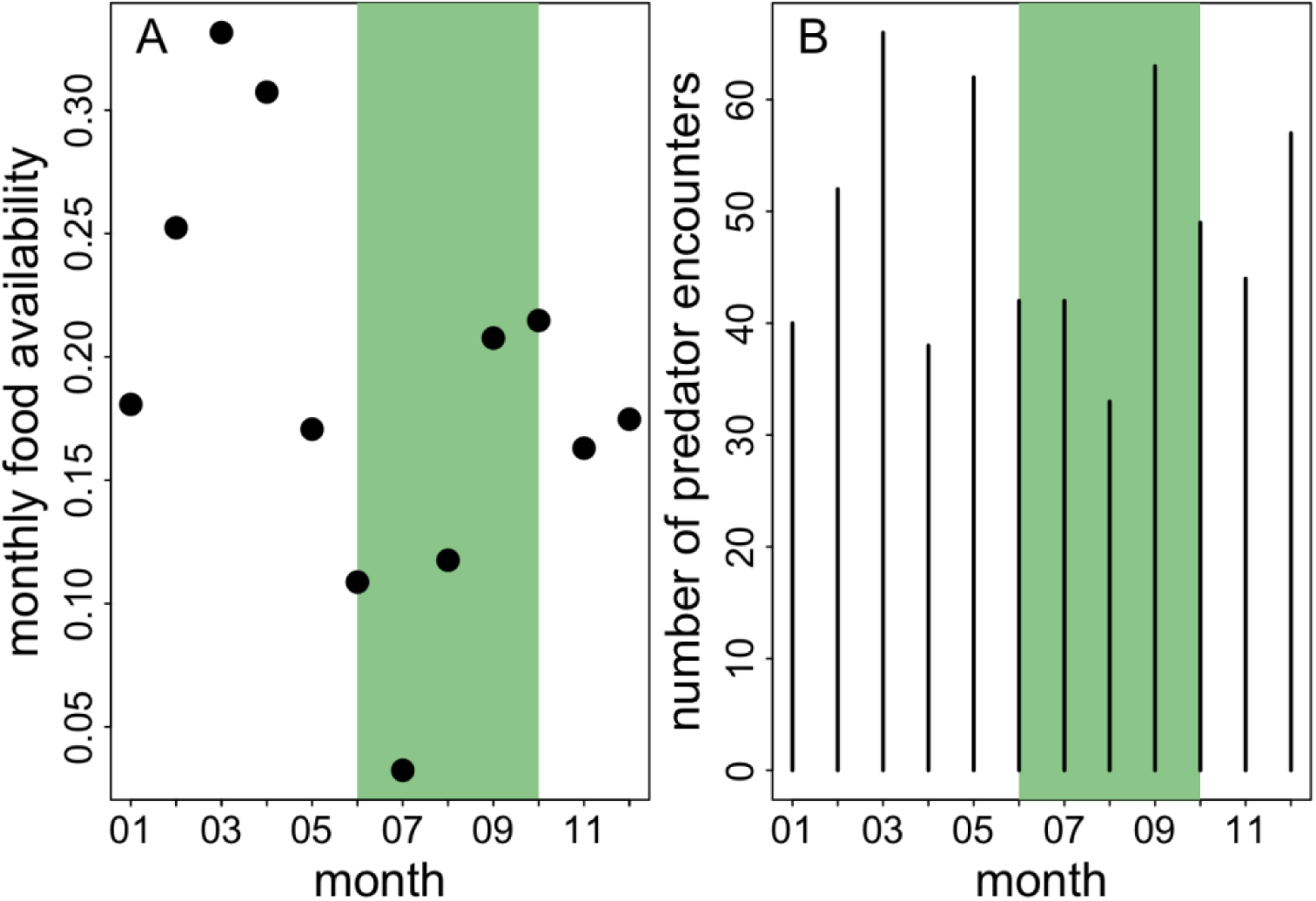
Monthly food availability scores for 2022 (A) and an overview of the number of predator encounters per month (B). The green area indicates the rainy season (June to mid-October).

Additionally, there was one sighting of a pack of eleven African wild dogs and four records that could not be reliably identified as either leopard or lion.

The average number of predator encounters varied depending on the time interval chosen to estimate predator presence. In 14-day intervals, we recorded 25 predator encounters on average (median, range 17 – 42). The average number of predator encounters in 7-day intervals was 13 (median, range 7 – 27). In 30-day intervals, the average number of predator encounters was 53 (median, range 39 – 82) and in 2-day intervals it was 3 (median, range 2 – 13).

### Distance to other parties

The minimum distance between sleeping sites of Guinea baboon parties in 2022 was 48.4 m on average (median, 16.8 – 136.7 m IQR) (Figure 7). We did not find evidence for an effect of food availability or predator presence on the minimum distance between sleeping sites of our study parties (Table 1, Figure 8). The result remained essentially the same for models using a 2-, 7- or 30-day time interval to assess predator presence (Tables S1-S3).

**Figure 7:**
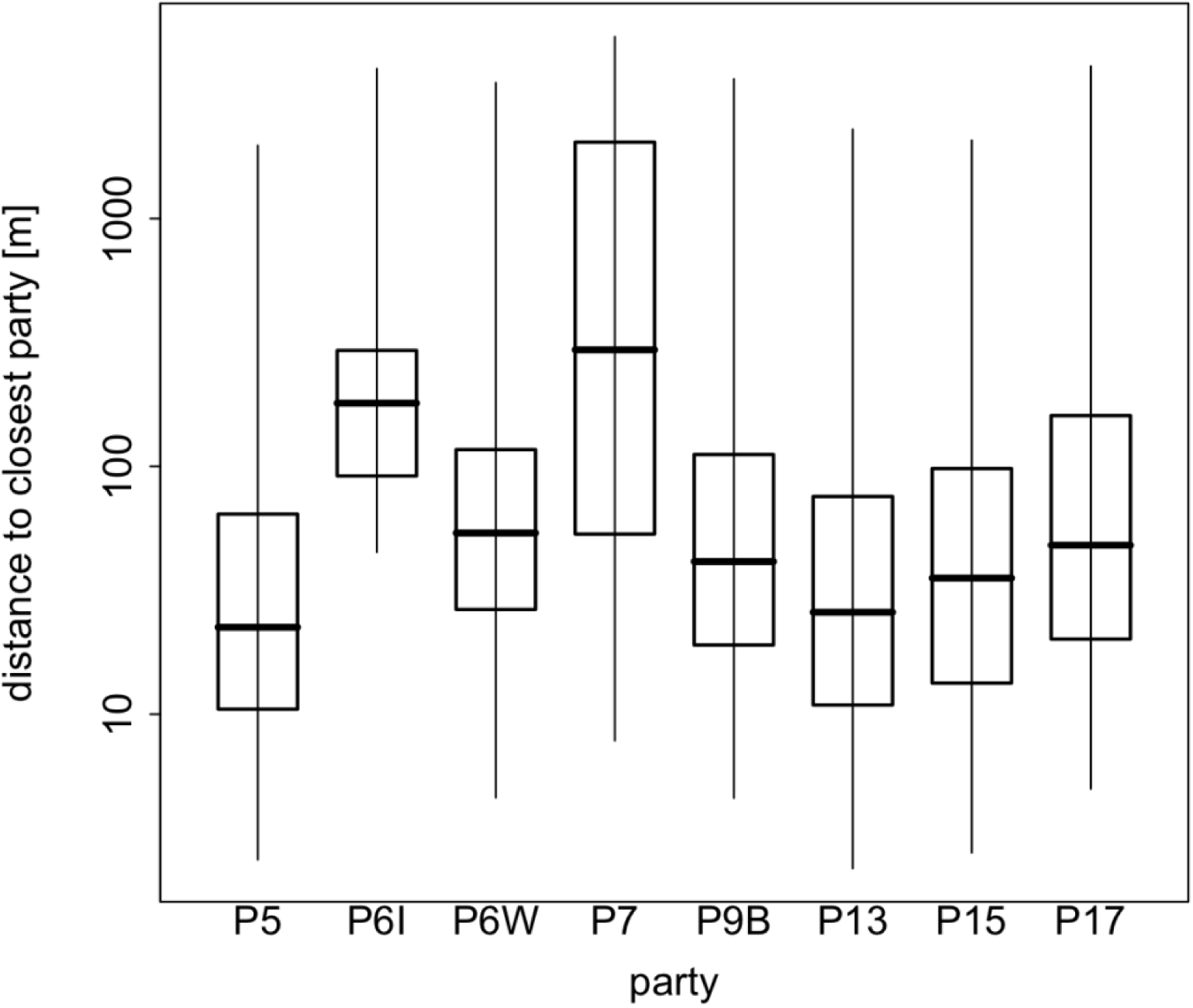
Minimum distances between sleeping sites of parties in 2022. Boxplots show the median (black line) and IQR with the lower (25%) and upper (75%) quartile. Whiskers represent the 2.5th and 97.5th percentiles. The distance to the closest party is depicted on a log scale for visual clarity.

**Figure 8:**
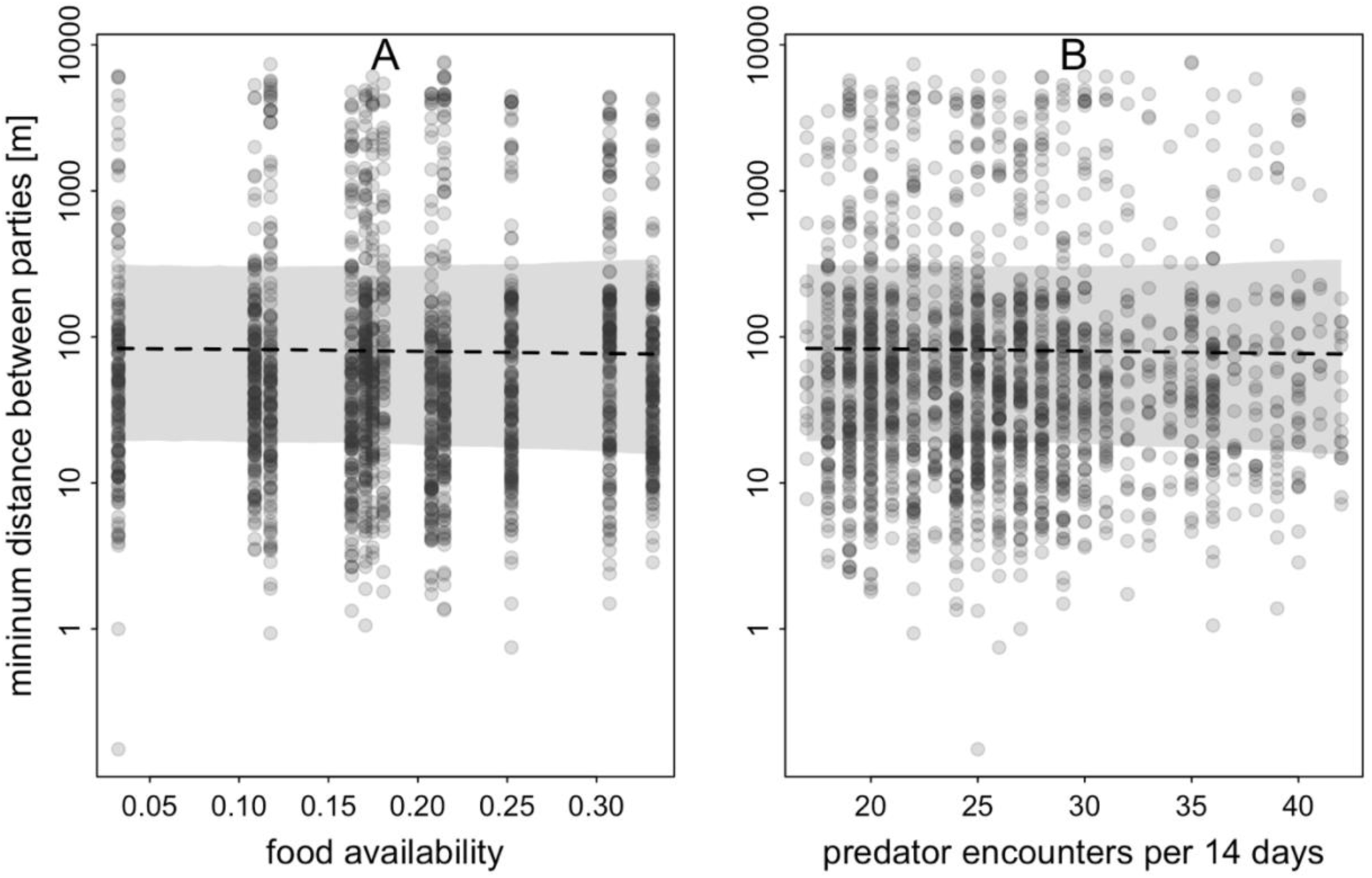
Model results for the effect of food availability (A) and the number of predator encounters per 14 days (B) on the minimum distance between sleeping sites of parties in 2022. For A, predator presence was centred to a mean of zero and for B, food availability was centred to a mean of zero. The minimum distance between parties is depicted on a log scale for visual clarity. The posterior mean is depicted as a dashed line and 95% credible intervals are shaded in grey.

**Table 1:**
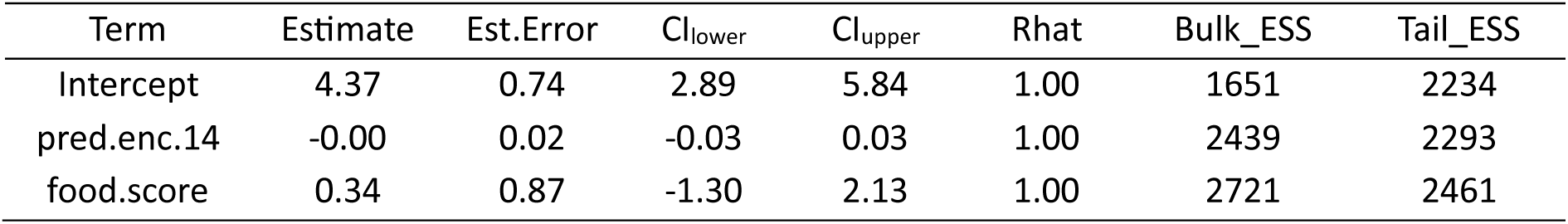
Model results for minimum distances between sleeping sites of parties in response to food availability and number of predator encounters within two weeks (estimates, standard errors, 95% credible intervals, Rhat, as well as Bulk and Tail Effective Sample Sizes).

### Patterns of sleeping site use

Guinea baboons used sleeping sites that were, on average, 2102 ± 1790 m (mean, SD, range 1 – 6237 m) from their previous sleeping site. Visual inspection of the distances between successively used sleeping sites for each party indicated that sites were rarely used continuously for extended periods (Figure 9). Comparing observed patterns of sleeping site use with random permutations of visit orders showed a tendency for more shorter and intermediate distances between sleeping sites than would be expected by chance, though the observed patterns were not markedly different from random expectations (Figure 10).

**Figure 9:**
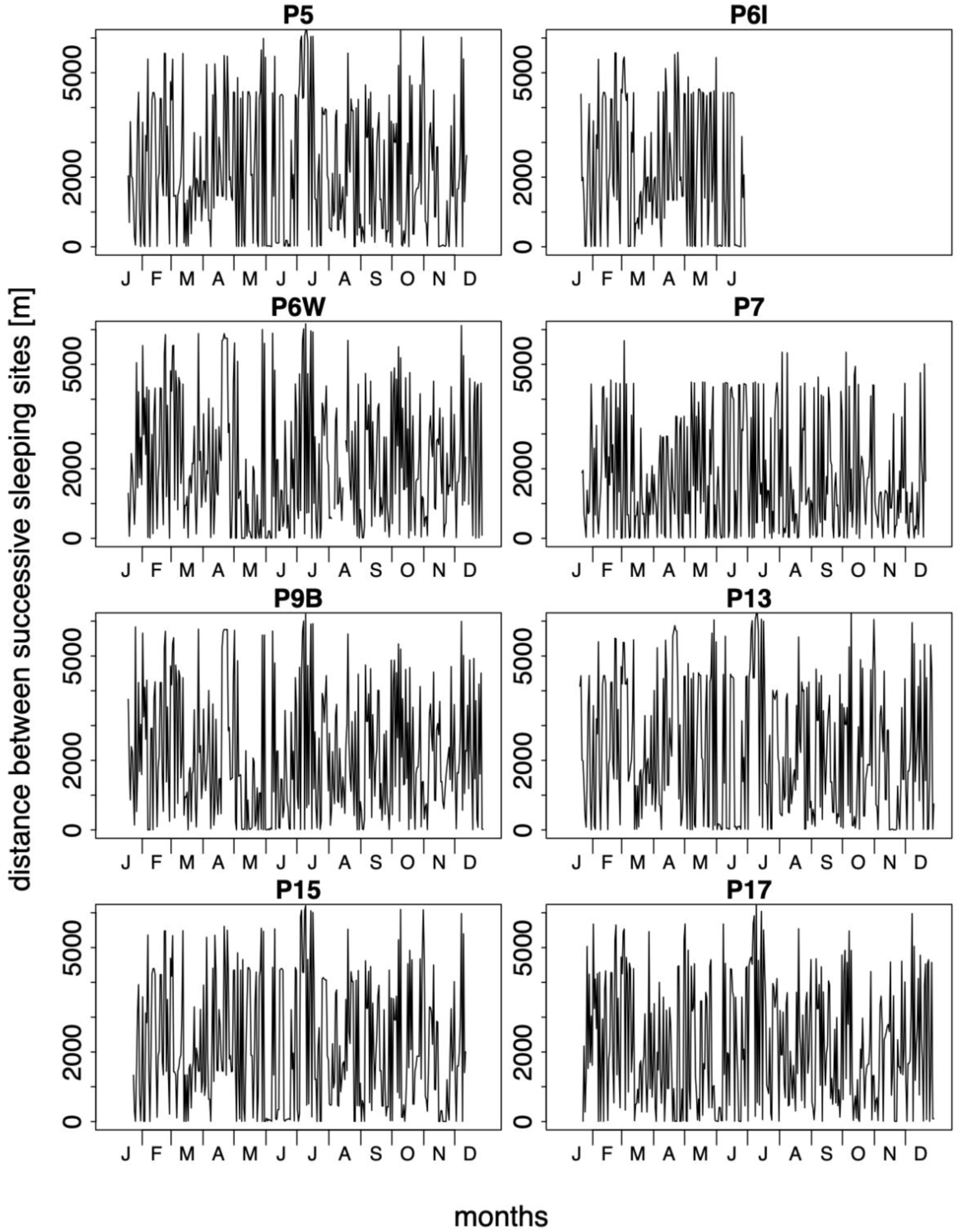
Distances between successive sleeping sites for all 2022 study parties across time. The collar for P6I failed in June 2022 due to water damage.

**Figure 10:**
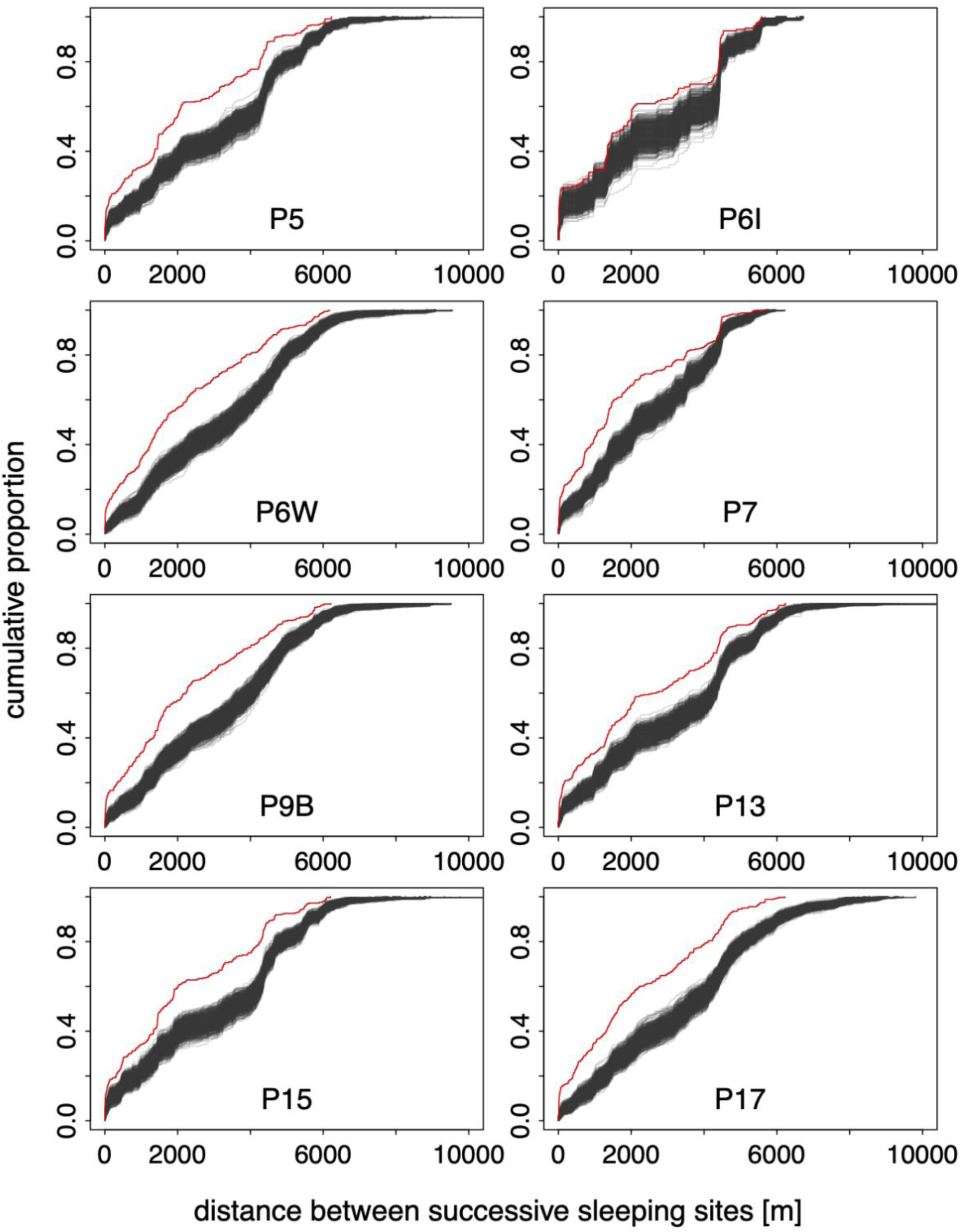
Patterns of sleeping site use of parties in 2022. Grey lines indicate the distance between successively used sleeping sites in 1000 randomised visit orders, and the red line represents the observed distances between successive sleeping sites.

## Discussion

### Sleeping sites and habitat types

Guinea baboons in Simenti spent most of their nights at sleeping sites in the gallery forest along the Gambia River. This frequent use of specific habitat types may reflect the availability of preferred sleeping tree species, such as *Ceiba pentandra*, *Celtis integrifolia*, or *Borassus akeassii*, which predominantly occur in habitats close to perennial water bodies. Trees of these species might offer better protection against nocturnal predation than other tree species due to their height (15 – 20 m) and smooth bark or thorns around the trunk, making them more difficult to climb for predators (Sharman, 1982; Anderson & McGrew, 1984; Zinner et al., 2021). Sleeping trees located in the gallery forest with a canopy cover of 75-100% (Klapproth, 2010) may also provide better connectivity to surrounding trees, facilitating an escape during a nocturnal attack (Harrison et al., 2020). However, *Borassus akeassii* is abundant throughout the study area.

The intensive use of sleeping sites in gallery forests, wetlands or temporary wetlands rather than in savannah habitats, where suitable sleeping trees such as *Borassus akeassii* are also abundant, may be associated with proximity to drinking water. During the dry season (November-May), the only available water sources in the study area are the Gambia River, one perennial wetland, and a few water bodies that usually dry out with the progression of the dry season. Sleeping near these water sources allows Guinea baboons to drink in the mornings and evenings. Similarly, Guinea baboons close to Mont Assirik in the south-eastern part of the Niokolo-Koba National Park chose sleeping sites that were 5 m from water bodies on average (Sharman, 1982).

Interestingly, in 2022, there was a shift in sleeping site use during the dry season, although this pattern was not evident in previous years. In 2022, Guinea baboons spent, on average, 16.4% (mean, range 3 – 29%) of their nights in sleeping sites located in savannah habitats (savannah woodland or tree/shrub savannah) compared to 0-3% during the rainy season. This shift might be due to feeding trees and herbaceous food items being available in savannah habitats during these months, with tree species like *Borassus akeassii*, *Strychnos spinosa*, *Piliostigma thonningii*, and *Acacia seyal* providing ripe fruit.

Our study parties may have used sleeping sites closer to these abundant food resources to minimise travel distances. Hence, time and energy were used towards approaches to such resources, a pattern seen in other primate species as well. Hamadryas baboons are thought to change their sleeping cliffs alongside changes in their foraging regions (Sigg & Stolba, 1981) and occasionally sleep on doum palm trees (*Hyphaene* spp.). In this way, the baboons likely have easier access to highly valuable palm fruits when these are scarce elsewhere in their habitat (Schreier & Swedell, 2008). However, the pattern we observed in 2022 was not evident in the other years of the study (2010-2012), suggesting that the variation in sleeping site use across habitats is likely driven more by opportunism than by successful resource exploitation and varies considerably between years, without any detectable consistent pattern.

### Distance to other parties

Guinea baboons almost always used sleeping sites close to at least one other party. The average minimum distance to the sleeping site of the closest party was less than 50 m, suggesting they may have even shared the same stand of trees. We did not, however, find any effect of food availability or predator presence on the distance between the sleeping sites of the closest parties. The lack of an observed effect of predator presence suggests that large aggregations at sleeping sites may not be related to communal predator detection or defence. However, it is essential to acknowledge that the absence of observed predators does not necessarily mean that predators were absent. Nocturnal predators such as leopards may have been nearby but undetected by the camera traps, observers, or the baboons themselves (Bidner et al., 2018).

Further, leopards preying on baboons at night do not seem to rely on surprise attacks (Busse, 1980), suggesting that larger groups do not benefit substantially from enhanced detection or mobbing behaviour at night. However, these groups might still benefit from the dilution of risk, by which the probability of any individual being targeted by a predator decreases as group size increases (van Schaik, 1983; Bidner et al., 2018). The formation of larger aggregations at sleeping sites (e.g., bands or gangs) may thus be primarily driven by the dilution of risk and social benefits. Larger aggregations at sleeping sites provide better opportunities for social interactions, offering more potential partners for socialising within and between parties. Guinea baboons often spend extended periods huddling, grooming, and greeting each other at sleeping sites before starting their morning foraging activities (Sharman, 1982; Patzelt et al., 2011).

### Patterns of sleeping site use

Upon visual inspection of the distances between successive sleeping sites, it seemed as if Guinea baboon parties tended to use sleeping sites that were either very close to, or more than 3 to 4 km away from, previously used sites, with fewer sites used at intermediate distances. When comparing the observed patterns to the expected distribution assuming random visit orders, we found that the observed patterns did not coincide with any of the scenarios we initially proposed. None of our study parties showed a pattern of sleeping site use consistent with either the predator avoidance hypothesis or minimising parasite load from faecal matter build-up. The only discernible trend was slightly more frequent use of sleeping sites that were closer to previously used ones than would be expected by chance. This result suggests that predator presence or parasite load are no significant factors influencing the Guinea baboons’ sleeping site use, at least not in the observed patterns. It is possible that other, less obvious factors are at play or that Guinea baboons in Simenti do not need to pay attention to these risks when selecting sleeping sites but instead use them opportunistically.

The parasite load hypothesis is inherently challenging to support or refute without detailed knowledge of the parasite communities present in the study area. Firstly, parasitological studies on Guinea baboons have reported a prevalence of gastrointestinal parasites of up to 78%, indicating that a significant portion of the population is already infected (N’da et al., 2022). Secondly, decay rates of faecal matter are largely unknown and may fluctuate substantially within and between years, as well as across different habitats. Finally, even if our study parties did not use a given sleeping site, other parties that inhabit the same area likely did. The population size of Guinea baboons in the Niokolo-Koba National Park is likely close to its carrying capacity, with estimates of 100,000 to 250,000 individuals (Galat-Luong et al., 2006; Rabeil et al., 2018). Given that Guinea baboons are likely exposed to faecal matter from conspecifics and other primates throughout their home ranges, for example, at feeding sites previously used by other parties, it seems unlikely that they would avoid these accumulations only at sleeping sites. In conclusion, the idea that they aim to reduce their parasite load by switching sleeping sites may be overly simplistic and not universally applicable.

### Limitations and prospects

It is important to acknowledge several methodological shortcomings of this study. We were unable to identify individual sleeping sites or assess their specific properties, occupancies and vacancies. Any attempt to delineate individual sleeping sites would have relied on arbitrary distance cut-offs, especially when looking at the almost complete use of gallery forests alongside the Gambia River as sleeping sites, particularly in the vicinity of the CRP Simenti campsite (Figure 2).

Therefore, our conclusions about sleeping site use are based on distances between successively used sleeping sites rather than on observed occupancy or vacancy patterns. Moreover, we were able to collar only a fraction of the baboon population present in the study area, suggesting that true patterns of sleeping site use may differ substantially from what we observed in our study parties. Sleeping sites might be used and reused at an even higher frequency than what we found in our study, which would offer even less support for the parasite avoidance hypothesis.

The lack of observed effects of food availability and predator presence in our models may partly result from the relatively coarse assessment of both, which may not have accurately captured changes in levels of competition or the perceived risk potentially affecting the sleeping site use of our study parties. Our assessment of food availability across the entire landscape likely did not appropriately capture the food-related drivers of sleeping site selection, if there were any. While landscape-level food availability may reflect competition between parties during their diurnal foraging activities in general, it is less suited to grasp the local distribution and abundance of resources potentially impacting sleeping site use. Future studies should incorporate direct observations of feeding behaviour and the abundance and distribution of food resources at sleeping sites, both in the evenings before the baboons recede into a sleeping tree as well as in the mornings as they leave.

Lastly, the patterns of sleeping site use we observed may not be solely determined by a given party’s independent choices but are likely influenced by the movement decisions of other parties. Despite the wide availability of sleeping trees, parties consistently stayed close at night. Rather than being driven purely by the availability of sleeping sites, this cohesion may result from how parties navigate their daily travel routes, the timing of their arrival at sleeping sites, and whether other parties are already present. The consistent proximity between sleeping sites of parties highlights the need to further explore the social and spatial processes influencing sleeping site use. Future research should examine how space-use and travel decisions during the day, particularly in the afternoons, shape sleeping site selection from a between-party perspective.

## Conclusion

Sleeping sites for Guinea baboons in Simenti are not a limited resource and the baboon parties did not compete for them, as reported for other baboon species. Frequently used sleeping tree species, particularly *Borassus akeassii*, were available in most habitats and could be found across the entire study area in large numbers. We did not find any patterns of sleeping site use that indicate predator avoidance or a reduction of parasite load at frequently used sites. We, therefore, conclude that the hypothesis of competition for sleeping sites and pronounced pressure from parasite infection is not supported by our findings for the Guinea baboons near Simenti. These baboons likely used the abundantly available sleeping sites opportunistically, rather than employing strategies aimed at predator avoidance or reducing parasite load. Future research should explore how social and spatial dynamics during the day shape sleeping site use from a between-party perspective.

## Declarations

### Ethics Approval

The collaring of the baboons was approved by the Ethics committee of the German Primate Center (document number E4-21).

### Availability of data and materials

GPS data from 2010 to 2012 can be found on Göttingen Research Online: https://doi.org/10.25625/IHEZUE (Zinner et al., 2021). GPS and phenological data, as well as data on predator presence from 2022 are available from the corresponding author upon request.

## Funding

This research was supported by the Deutsche Forschungsgemeinschaft (DFG, German Research Foundation), Grant/Award Number: 254142454 / GRK 2070. This publication was supported by the Leibniz Association through funding for the Leibniz ScienceCampus Primate Cognition (W45/2019 – Strategische Vernetzung).

### Authors’ contributions

**Lisa Ohrndorf:** Conceptualisation (equal); investigation (lead); data curation (equal); formal analysis (supporting); visualisation (equal); writing – original draft (lead); writing – review and editing (equal). **Roger Mundry:** Data curation (equal); formal analysis (lead); visualisation

(equal); writing – original draft (supporting); writing – review and editing (equal). **Jörg Beckmann:** Investigation (supporting); writing – original draft (supporting); writing – review and editing (equal). **Julia Fischer:** Conceptualisation (equal); writing – review and editing (equal). **Dietmar Zinner:** Conceptualisation (equal); writing – original draft (supporting); Writing – review and editing (equal).

## Acknowledgements

We thank the Direction des Parcs Nationaux (DPN) and the Ministère de l’Environnement et de la Protéction de la Nature (MEPN) de la République du Sénégal for the approval to conduct this study in the Niokolo-Koba National Park. We particularly appreciated the support and cooperation of the former Conservateur of the park Jacques Gomis. We are grateful to all the CRP Simenti staff and field assistants for their support in the field, in particular Djibril Coly, Chérif Younousse Kéba Camara, and Amadou Bamba Diedhiou. We owe special thanks to Irene Gutiérrez Díez for her help with the data collection in the field.

