## Supplementary material for "Patterns of sleeping site use of Guinea baboon parties (*Papio papio*)"

Table S1: Model results on minimum distances between sleeping sites of parties in response to food availability and number of predator encounters within 7 days (estimates, standard errors, credible intervals, Rhat, as well as Bulk and Tail Effective Sample Sizes).

| Term | Estimate | Est.Error | $CI_{lower}$ | $CI_{upper}$ | Rhat | Bulk_ESS | Tail_ESS |
|---|---|---|---|---|---|---|---|
| Intercept | 4.45 | 0.64 | 3.19 | 5.73 | 1.00 | 1282 | 1954 |
| pred.enc.7 | -0.01 | 0.02 | -0.05 | 0.02 | 1.00 | 3140 | 2407 |
| food.score | 0.40 | 0.83 | -1.22 | 2.17 | 1.00 | 3051 | 2361 |

Table S2: Model results on minimum distances between sleeping sites of parties in response to food availability and number of predator encounters within 30 days (estimates, standard errors, credible intervals, Rhat, as well as Bulk and Tail Effective Sample Sizes).

| Term | Estimate | Est.Error | $CI_{lower}$ | $CI_{upper}$ | Rhat | Bulk_ESS | Tail_ESS |
|---|---|---|---|---|---|---|---|
| Intercept | 4.60 | 0.76 | 3.15 | 6.14 | 1.00 | 1275 | 2063 |
| pred.enc.30 | -0.01 | 0.01 | -0.02 | 0.01 | 1.00 | 1989 | 2217 |
| food.score | 0.41 | 0.89 | -1.29 | 2.25 | 1.00 | 2411 | 2393 |

Table S3: Model results on minimum distances between sleeping sites of parties in response to food availability and number of predator encounters within 2 days (estimates, standard errors, credible intervals, Rhat, as well as Bulk and Tail Effective Sample Sizes).

| Term | Estimate | Est.Error | $CI_{lower}$ | $CI_{upper}$ | Rhat | Bulk_ESS | Tail_ESS |
|---|---|---|---|---|---|---|---|
| Intercept | 4.30 | 0.57 | 3.15 | 5.43 | 1.00 | 1498 | 2175 |
| pred.enc.2 | 0.02 | 0.04 | -0.07 | 0.11 | 1.00 | 2226 | 1939 |
| food.score | 0.19 | 0.81 | -1.34 | 1.87 | 1.00 | 2499 | 2325 |

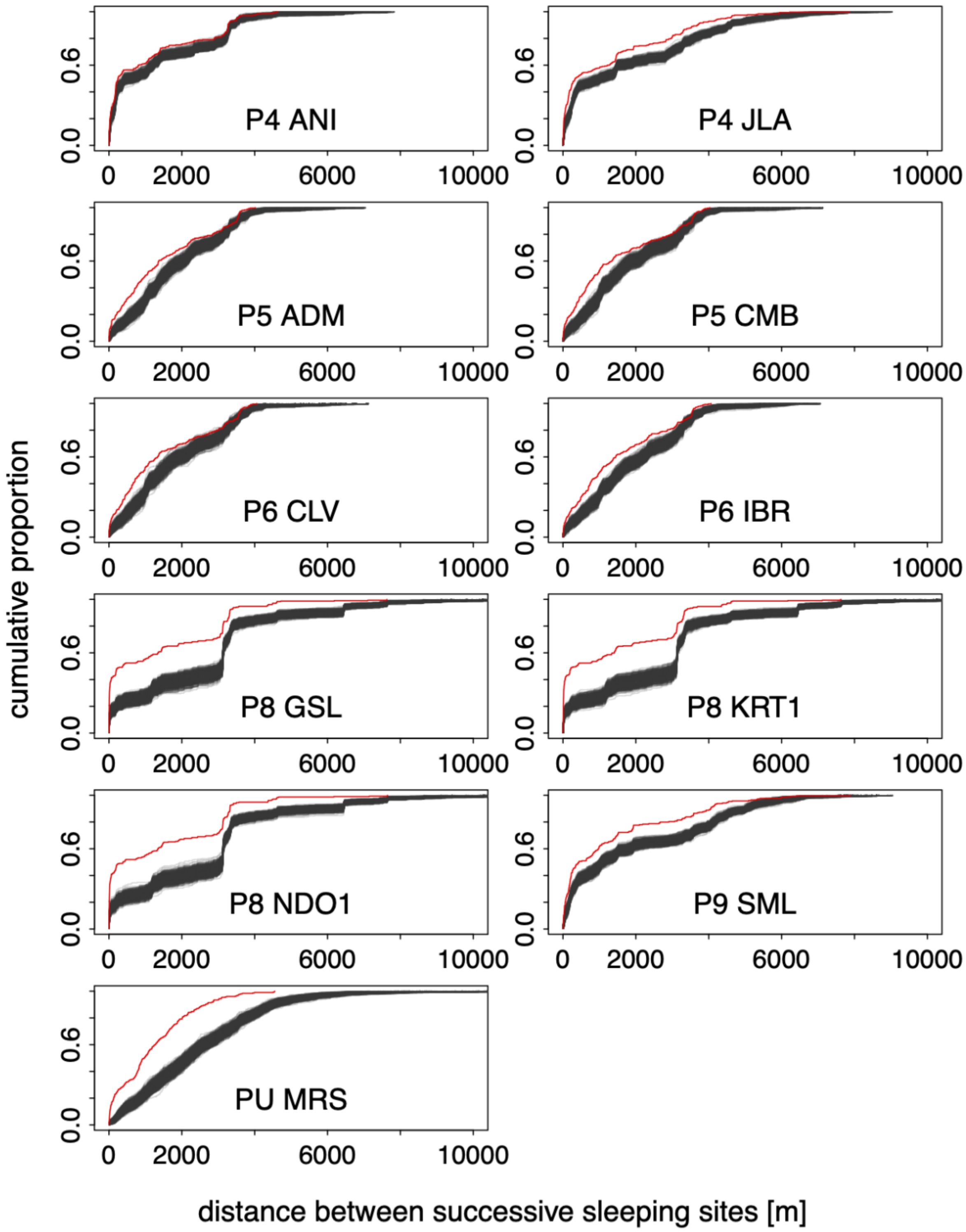


Figure S1: Patterns of sleeping site use of parties and associated collared individuals in 2010. Grey lines indicate 1000 randomised visit orders, and the red line depicts observed distances between successive sleeping sites.

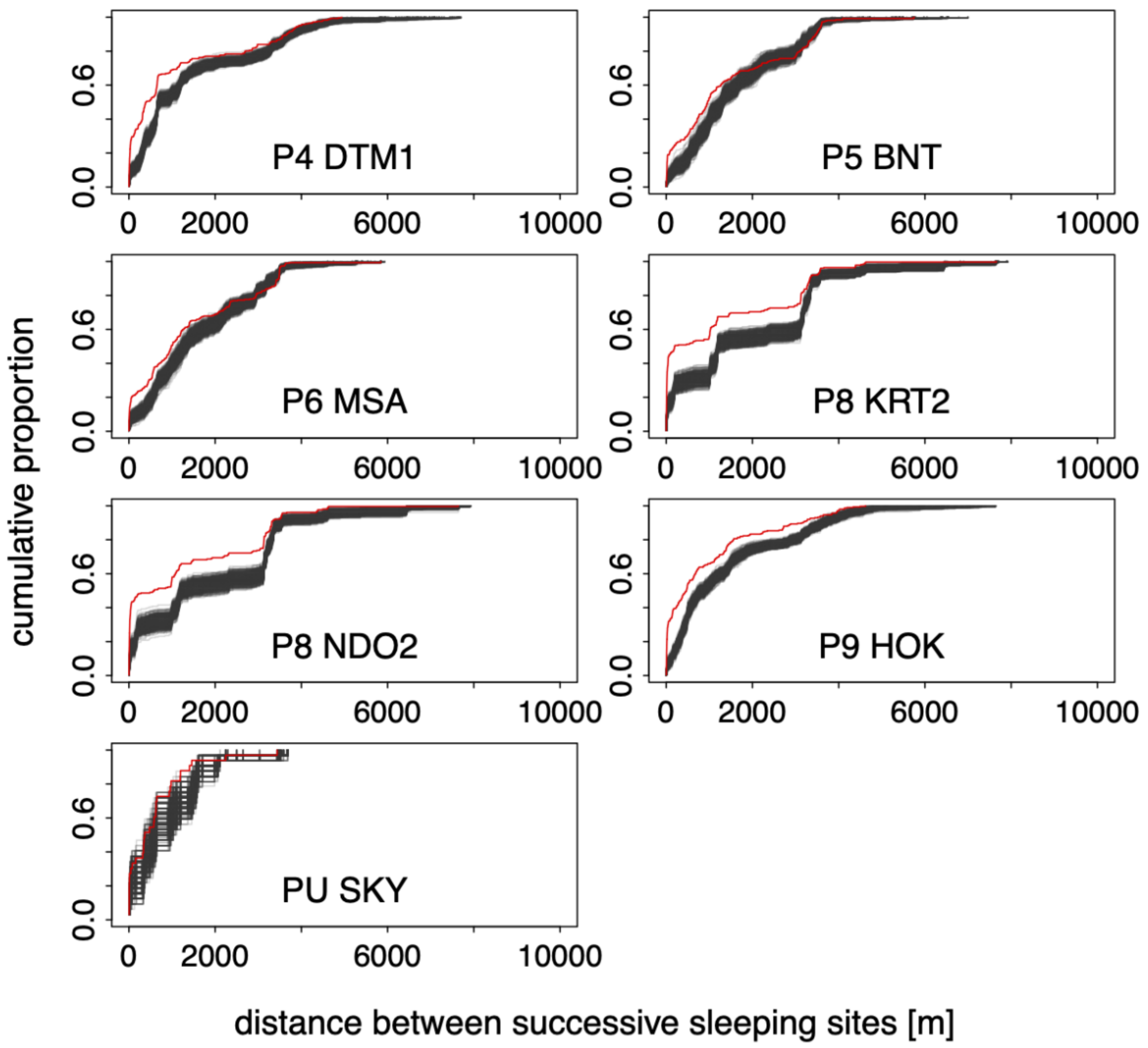


Figure S2: Patterns of sleeping site use of parties and associated collared individuals in 2011. Grey lines indicate 1000 randomised visit orders, and the red line depicts observed distances between successive night sleeping sites.

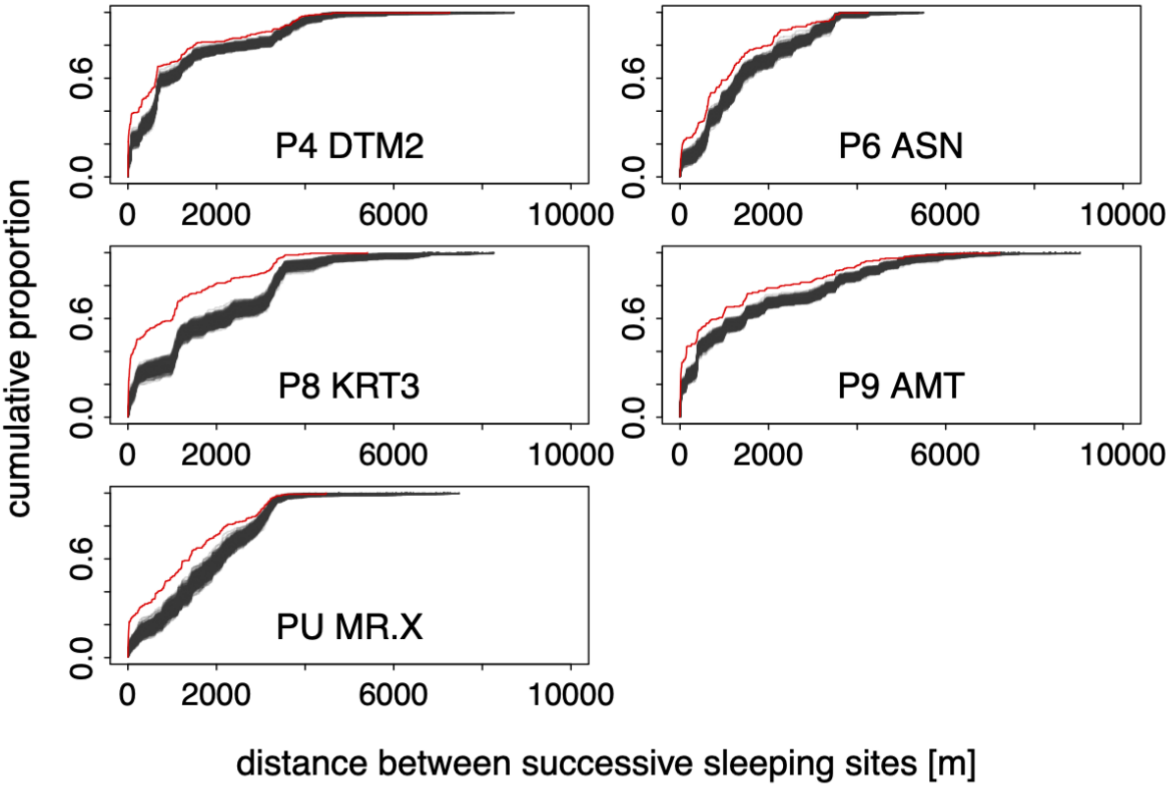


Figure S3: Patterns of sleeping site use of parties and associated collared individuals in 2012. Grey lines indicate 1000 randomised visit orders, and the red line depicts observed distances between successive sleeping sites.
